## Supplementary Information for "Superresolved microparticle traction force microscopy reveals subcellular force patterns in immune cell-target interactions"

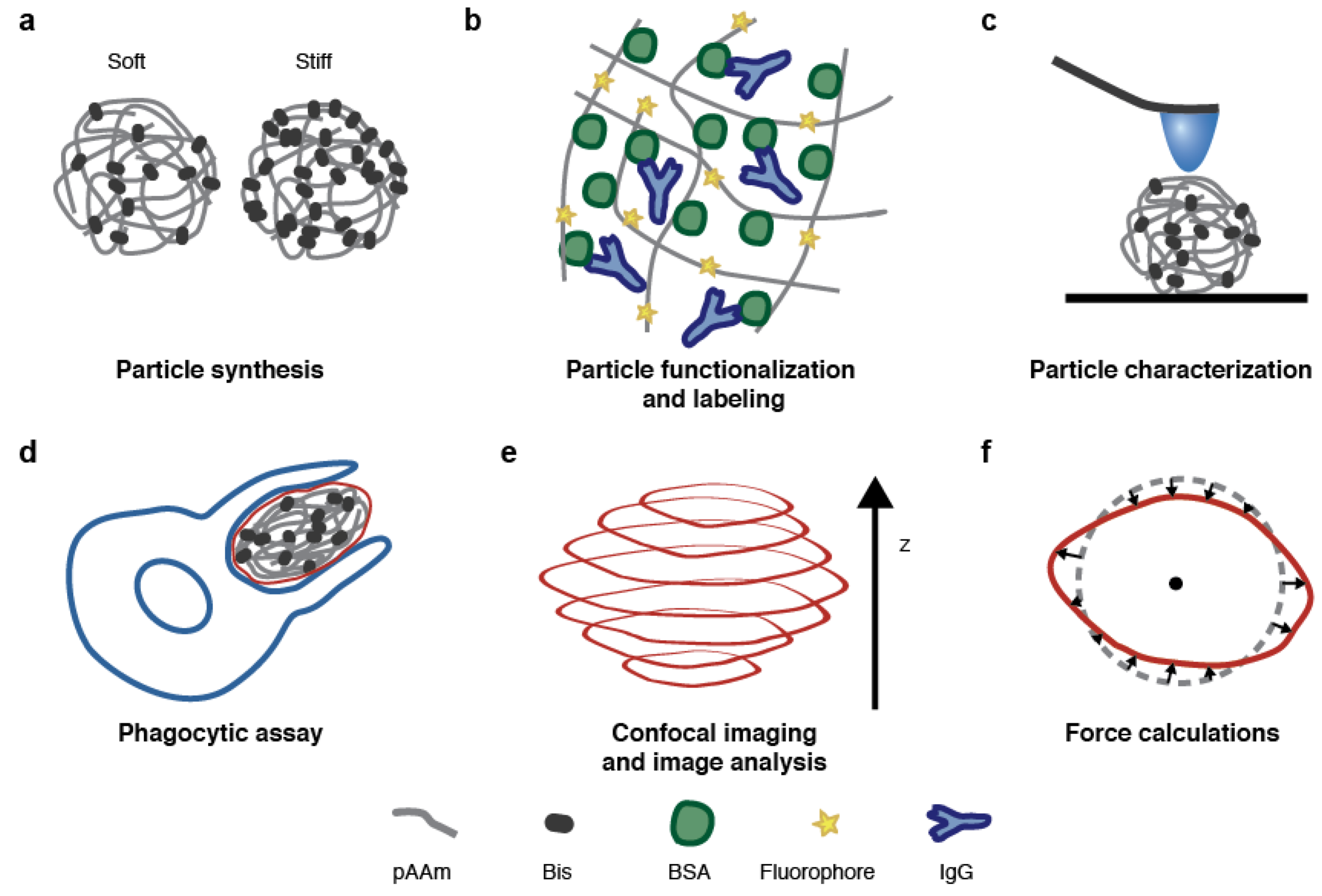

**Supplementary Figure 1: Schematic diagram of the developed method. a,** Rigidity of spherical deformable polyacrylamide-co-acrylic acid microparticles (DAAM-particles) can be tuned by adjusting the crosslinker (bisacrylamide) or total acrylamide concentration. **b,** Inclusion of acrylic acid (AAc) in the DAAM-particle allows direct conjugation of proteins and carboxyl-reactive fluorescent dyes to the gel. In our approach, the gel is functionalized further with αBSA-IgG. **c**, The mechanical properties of DAAM-particles can be characterized using atomic force microscopy (AFM). Using pyramidal tips with large end radius (~700 nm) allows both imaging and Hertzian indentations of MPs. **d,** Phagocytes (in this manuscript J774 macrophage-like cells) are exposed to DAAM-particles functionalized to trigger phagocytosis. When the DAAM-particles are of sufficiently low rigidity, the forces exerted by the cell during phagocytosis will deform the particles. **e,** Confocal microscopy allows visualization of the particle shape using the dye embedded in the gel. Image analysis allows reconstruction of the particle boundary with superresolution in the radial direction. **f,** As the mechanical properties of the DAAM-particles are known, isotropic and homogeneous, observed deformations can be used to calculate the forces exerted by the cell.

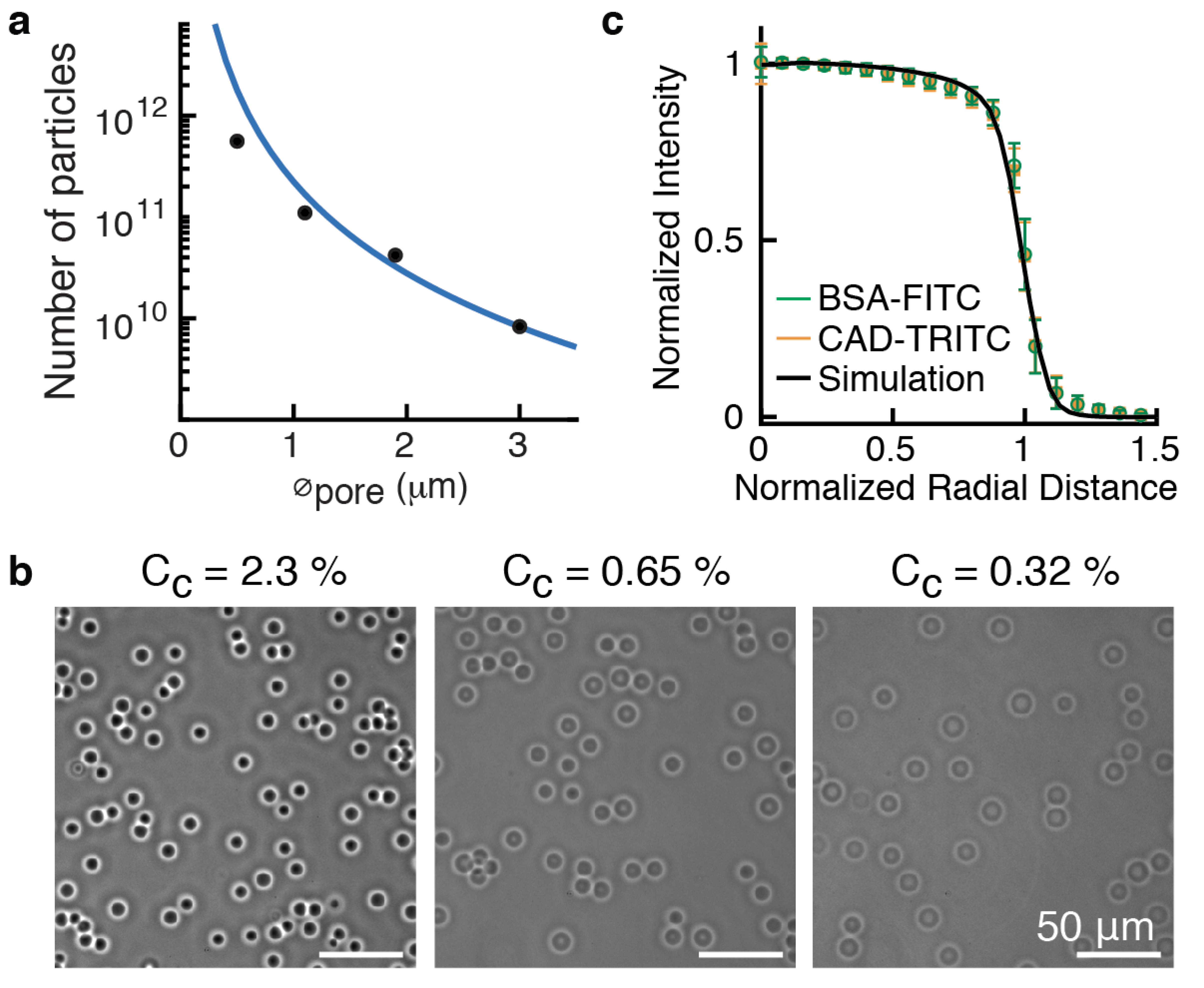

**Supplementary Figure 2: Batch production of pAAm Microparticles using hydrophobic Shirasu Porous Glass (SPG). a,** SPG membrane pore size versus estimated number of particles per batch, calculated as V_aq_/V_MP_, where V_aq_ is the volume of the aqueous phase containing the gel mixture (10 mL), and V_MP­_ is the volume of a single particle, as quantified in figure 1b (main text). The blue line corresponds to the linear relation between SPG membrane pore size and particle size with slope 4.9 (Fig. 1b, main text). Note that this estimate assumes no gel swelling, which results in an underestimate of the number of particles per batch. **b,** Phase images of MPs that were made with the same pore size membrane and different crosslinker concentrations (C_c_). Lowering C_c_ results in more hydrogel swelling (as evidenced from the reduced contrast indicating lower polymer density) and larger particles. **c,** Radial fluorescence intensity profile illustrating particle radial homogeneity (more than 100 MPs were used to create this average). Identical to figure 1e (main text), but for C_c_ 0.32%.

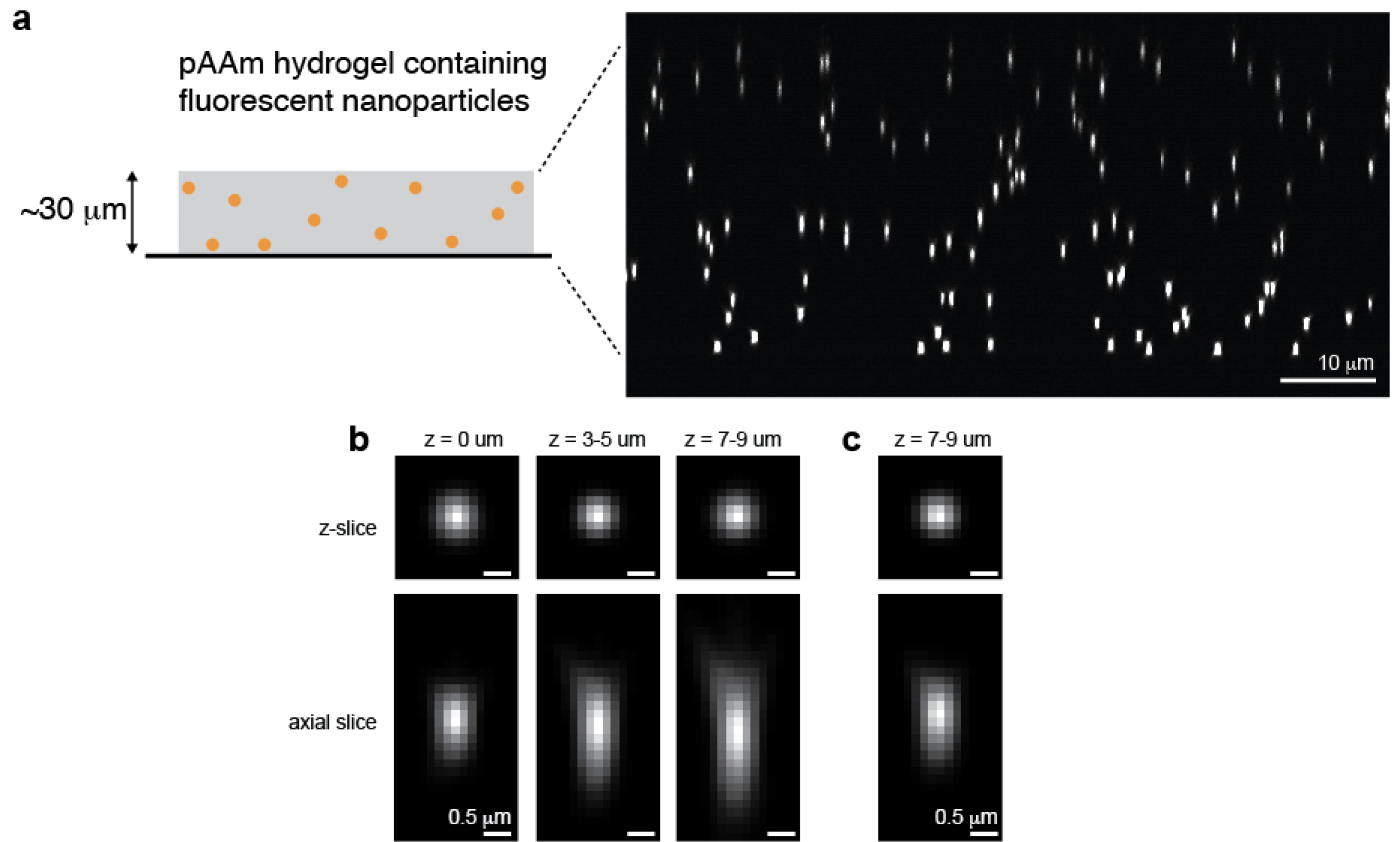

**Supplementary Figure 3: Depth-dependent measurement of point-spread-functions (PSFs) in adherent hydrogels. a,** (Left) Diagram of the approach: polyacrylamide (pAAm) hydrogels that are covalently bound to the glass surface, and, in which fluorescent nanoparticles (~100 nm diameter) were incorporated, were imaged on a confocal microscope. (Right) Confocal image (axial-slice) of such a hydrogel. It can be seen that particles further from the surface appear dimmer and elongated along the axial direction. The location of the glass slide can be determined from the background signal, when the contrast is enhanced (now shown). **b,** Reconstructed point-spread-functions (PSFs) from red 100 nm nanoparticles imaged at various distance from the surface (z). Fluorescent signals of over 50 nanoparticles within each distance interval were superimposed with subpixel resolution and averaged. In z-slices the particles look identical, but in axial-direction the PSF width increases with distance to the surface. The middle PSF was used for simulating the appearance of a homogeneous microsphere on the confocal (Fig. 1c, main text and Supplementary Fig. 2). The PSF on the right was used for deconvolution of DAAM-particles in PBS (Fig. 4, main text). **c,** Similar to **b,** but reconstructed from 100 nm yellow-green nanoparticles measured in high refractive index (RI) medium (Vectashield (VS); RI ~ 1.45). This PSF was used for deconvolution of DAAM-particles imaged in VS (Fig. 3, main text).

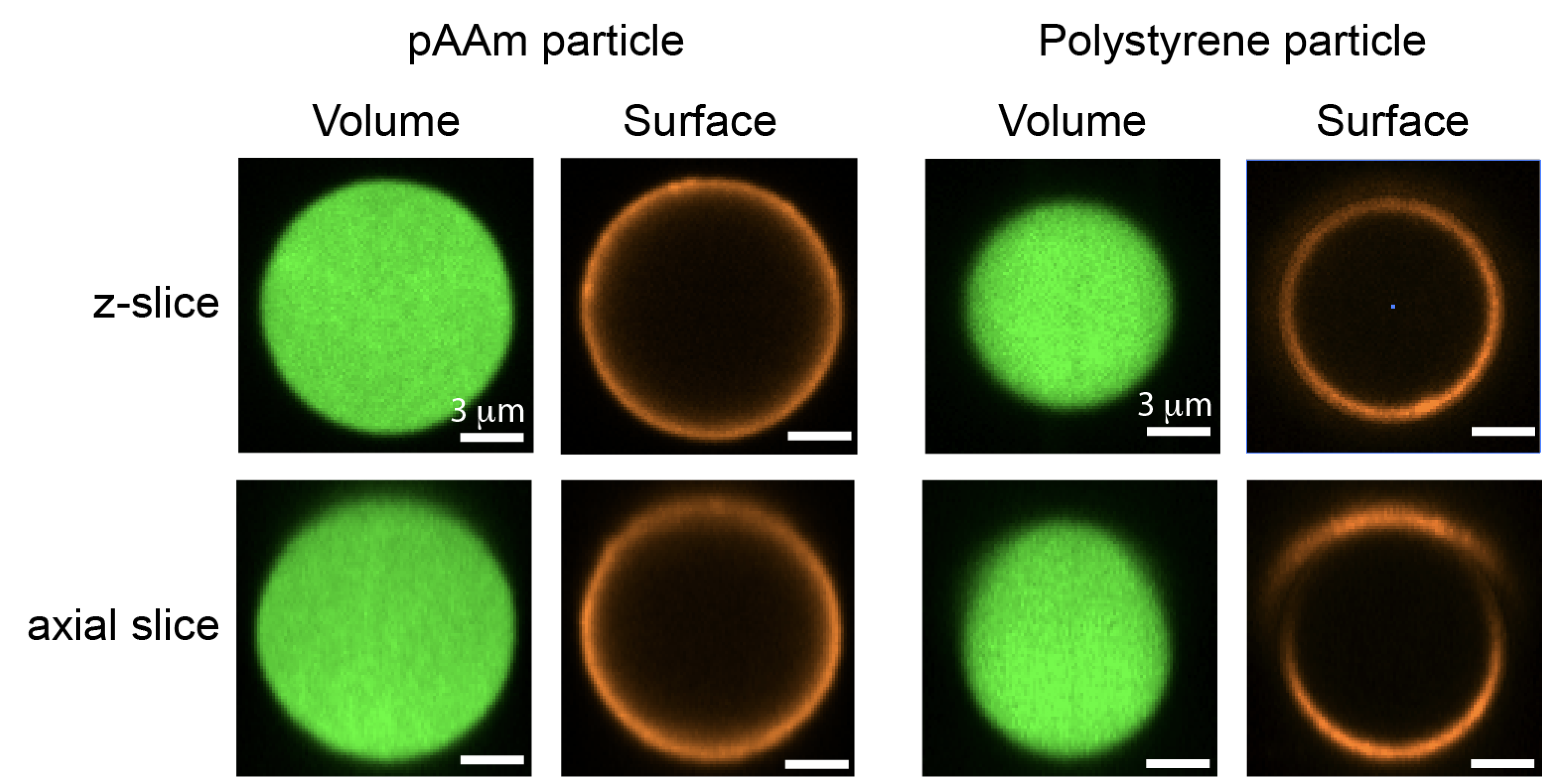

**Supplementary Figure 4: Microparticle lensing and image distortion in high refractive index media.** Z- and axial slices from representative confocal image stacks through a pAAm (C_c_ = 0.32%) and a 10 μm Yellow-Green PS particle. pAAm MPs were functionalized with Alexa 488-cadaverine. Surface of pAAm and PS MPs was functionalized with BSA, αBSA-IgG, and Alexa 546-GαR-IgG measured in Vectashield (refractive index: ~1.45). The polystyrene particle appears deformed (‘egg-shaped’) in the axial direction. The surface stain is also clearly distorted: the signal is displaced from the actual surface, and shows large intensity variations, in the upper hemisphere. These artifacts arise due to the refractive index mismatch between particle and medium, which results in the particle acting as a lens. Such gross artifacts are not noticeable with pAAm particles, since their refractive index is very close to the medium (Fig. 1d, main text).

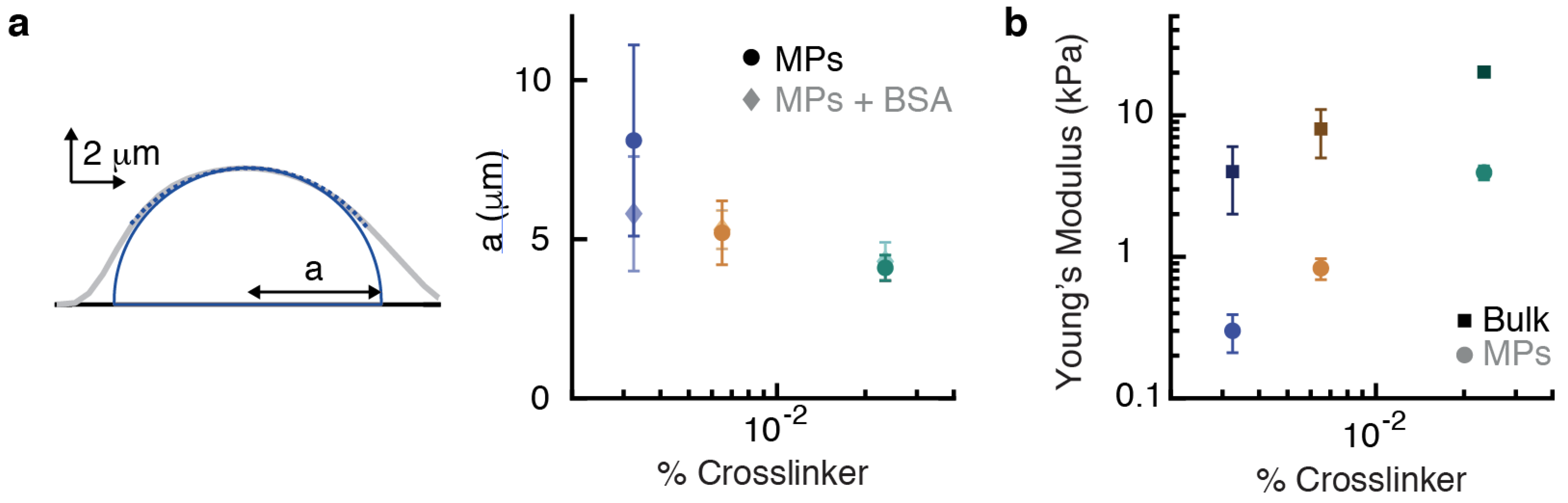

**Supplementary Figure 5: Contact radius determination and comparison of elastic properties between pAAm microparticles and bulk gels. a,** (Left) Shape determination from atomic force microscopy (AFM) images. In grey, a line profile from an image through the maximum of a particle, measured along the slow scanning axis. A spherical arc was fit to the upper half of the line profile (since the lower part of a sphere cannot be imaged with AFM). The particle radius of curvature (R) was then estimated as R = R_tot_ - R_tip_, where R_tot_ is the radius of the fitted spherical arc, and R_tip_ is the radius of the AFM tip. Under assumption that the particles form spherical caps, and using the height estimated from the image, this yields an estimate of the particle shape and, hence, the contact radius with the surface (a). (Right) Quantification of the particle contact radius with the surface for particles with and without functionalization, and various crosslinker concentration. Errorbars indicated standard deviations. **b,** Comparison of Young’s moduli of microparticles (~10 μm) with bulk adherent hydrogels of identical composition, *i.e.* same total acrylamide concentration, same crosslinker concentration and same acrylic acid concentration. n = 15 indentations in unique locations for all the bulk gels. Data for MPs is identical to figure 1f in the main text. Error bars mark standard deviations between individual force curves (bulk gels), or standard deviations from particle-to-particle, where for each particle the Young’s modulus was estimated as the average from 2-5 indentation curves (MPs).

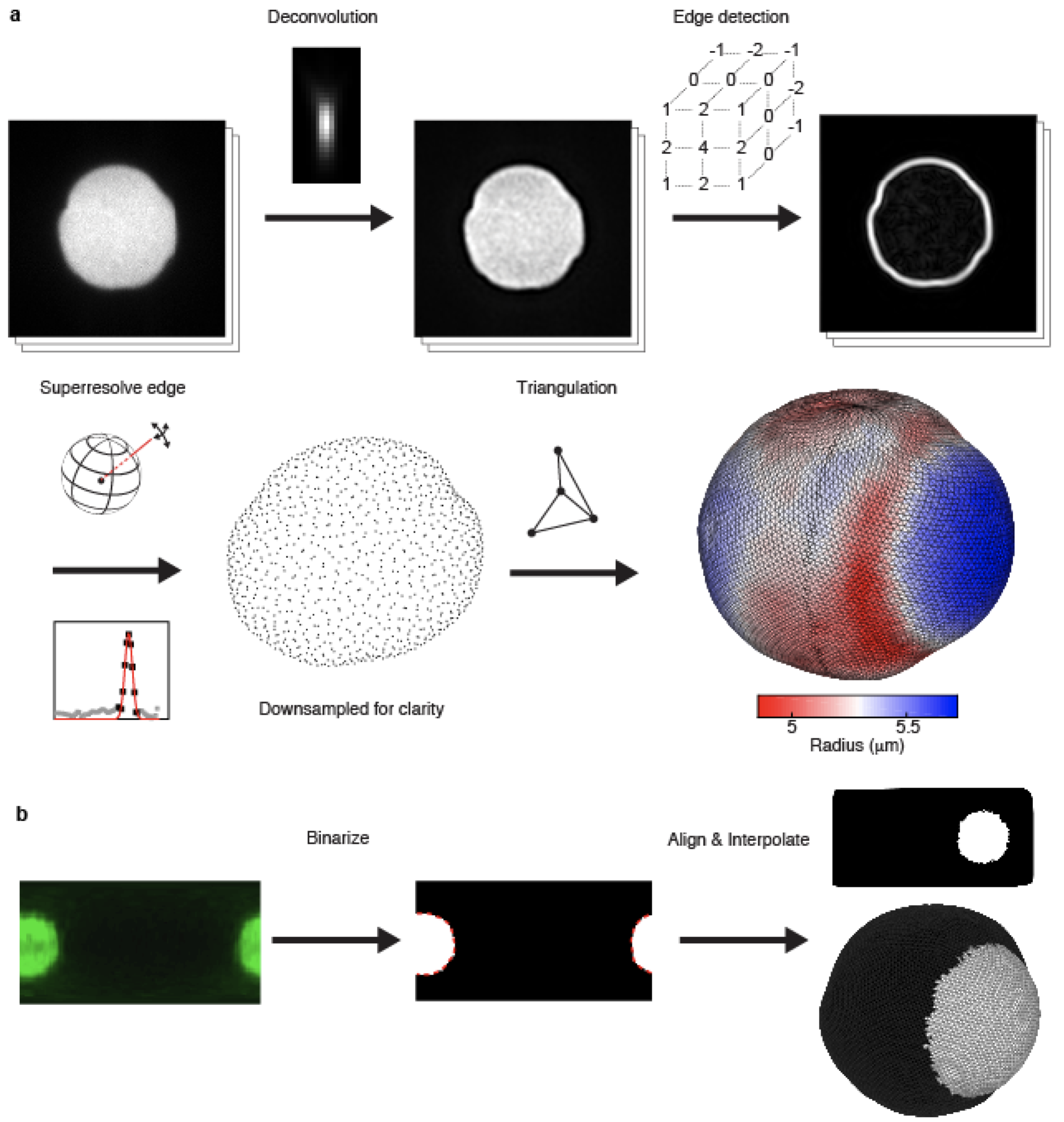

**Supplementary Figure 6: Image analysis steps for 3D particle shape reconstruction and free boundary analysis. a,** Particle shape reconstruction. Confocal image stacks were first deconvolved, using a point spread function measured with 100 nm red fluorescent nanoparticles, imaged ~8 μm from the surface (Supplementary Fig. 3). Subsequently, edge detection was performed using convolution with the 3 kernels of the 3D-Sobel Operator, after which a single 3D image was calculated as the magnitude of the 3 resulting images. Next, lines were drawn outward from the centroid with approximately equally spaced angles, and the fluorescent signal along these lines was approximated by interpolation. A Gaussian function was fit to the peak in the resulting line profile, allowing determination of surface coordinates. Finally, triangulation was performed to calculate particle and surface properties (*e.g.* centroid, volume, surface area, curvature). **b,** Free boundary analysis. The immunostained non-occluded part of the particle surface was analyzed similar to the particle fluorescent signal, except that 1) Edge detection was not performed; 2) A regular grid was used; 3) The peak of the Gaussian profile was not localized, but instead, the integral of the Gaussian was calculated, which was approximated as I_tot_ = 1.065 * I_max_ * FWHM, where I_tot_ is the total fluorescent intensity under the Gaussian, I_max_ is the maximum fluorescent intensity, and FWHM is the full width half maximum of the Gaussian. I_tot_ was more uniform over the MP surface than the I_max_. A 2D projection of I_max_ was made (left) and the signal was binarized using a region-based active contour (or “snake”) algorithm^1^ (dashed red line). Finally, the binary mask was used to align the particles, with the base of the cup at longitude -0.5π and latitude 0. The mask was also interpolated to determine for each edge coordinate if it was within the area covered by the cell, or part of the traction-free boundary.

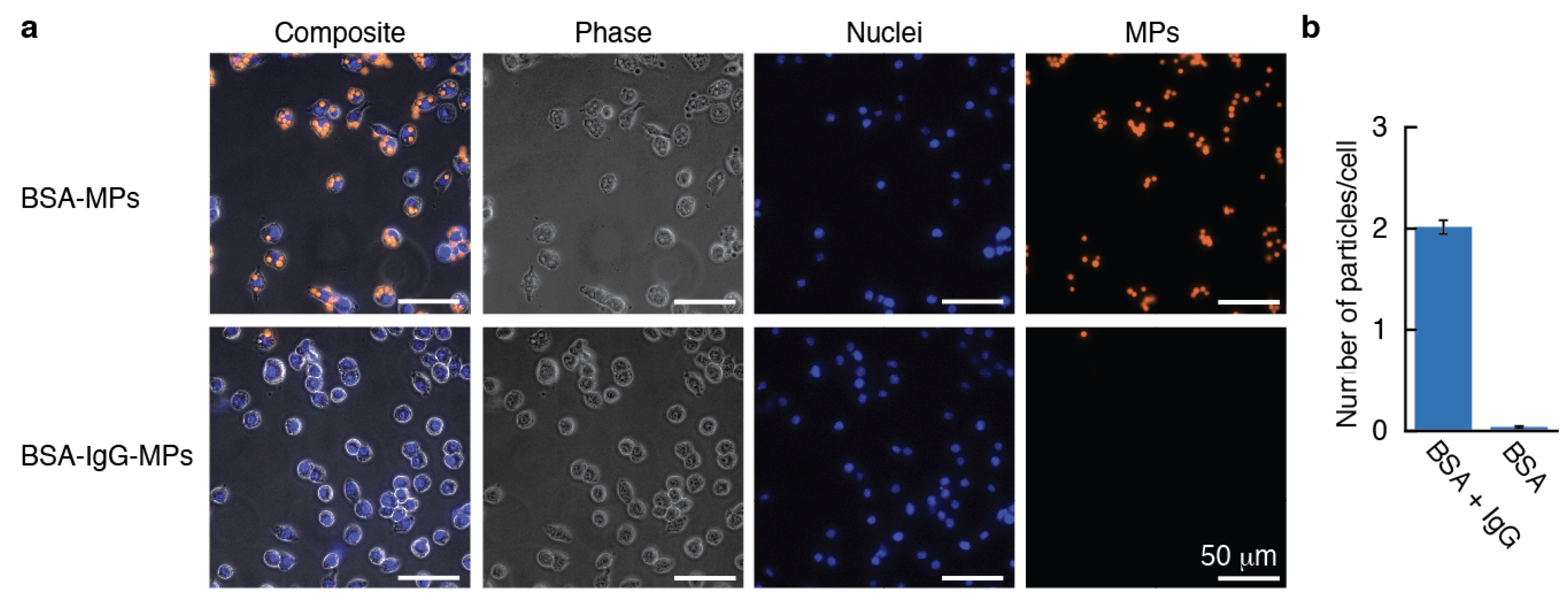

**Supplementary Figure 7: Uptake of pAAm MPs by J774 macrophage-like cells is ligand-dependent. a,** J774 murine macrophage-like cells were seeded in a 12-well plate (1.5 x 10^5^ cells/well). The next day, cell nuclei were stained with Hoechst, and then exposed to stiff (crosslinker concentration: 3.2%) MPs (~7.5 x 10^5 MPs/well), stained with TRITC-Cadaverine and further functionalized with BSA and αBSA-IgG, or only BSA, for 30 minutes. Cells were then fixed, washed vigorously and imaged (40x). Typical fields of view (FOV) are shown. **b,** Quantification of the number of beads (counted from fluorescent TRITC-CAD signal) per cell (counted from fluorescent Hoechst signal). Approximately 400 cells were analyzed in each condition. Error bars represent standard deviations estimated as √λ, where λ is the average number of beads per cell, hence we make the assumption that the number of MPs per cell follows a Poisson distribution.

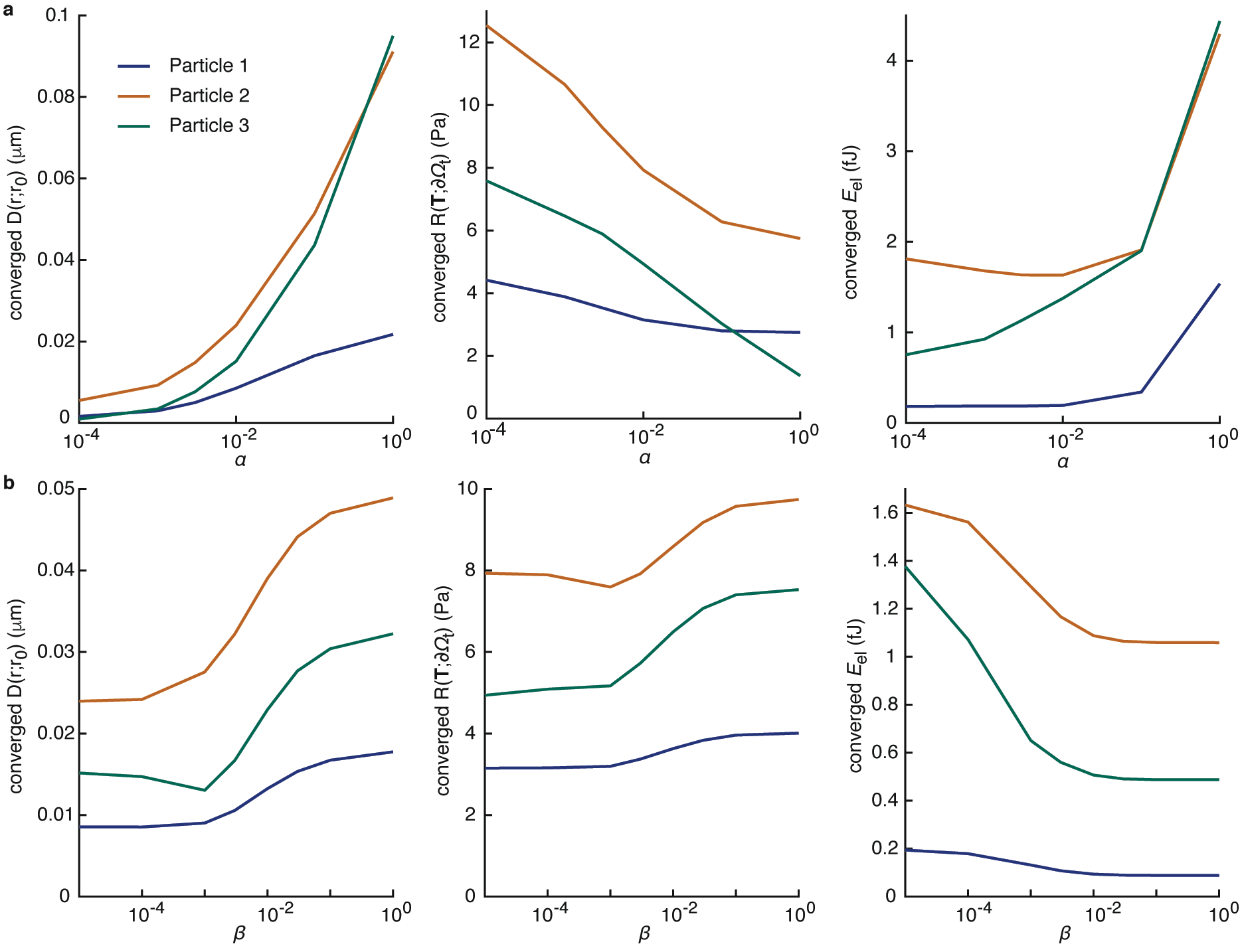

**Supplementary Figure 8: Determination of the free parameters αandβ in solving the mixed-boundary problem.** The converged shape difference $D\left( r;r_{0} \right)$, traction magnitude $R\left( \boldsymbol{T};\partial\Omega_{t} \right)$ on the traction-free region, and the elastic energy $E_{el}$ as a function of optimization parameters α and β for particle 1-3 (see main text). Data for particle 4 is not shown because it is fully internalized and therefore does not have a traction-free boundary. Note that the dependence on α and β is similar for all 3 particles with strongly different sizes of the traction-free boundary (9 - 84% of the total particle surface). This allows use of a single α and β value for all shapes. **a,** Converged $D\left( r;r_{0} \right), R\left( \boldsymbol{T};\partial\Omega_{t} \right)$ and $E_{el}$ as a function of α with $\beta$ ^-5^. $\alpha$ controls emphasis on the traction-free boundary during optimization: the higher the $\alpha$ value, the lower the traction magnitude will be on the traction-free region. At $\alpha\approx{10}^{-2}$, the converged shape difference (< 25 nm) and traction magnitude (< 10 Pa) are both low and the energy has plateaued. Increasing $\alpha$ leads to a large increase in the elastic energy, whereas reducing $\alpha$ increases $R\left( \boldsymbol{T};\partial\Omega_{t} \right)$ (it decreases $D\left( r;r_{0} \right)$, but the shape is already fitted below the noise level: ~50 nm). Therefore, $\alpha\approx{10}^{-2}$ is the optimal choice. **b,** plot of the converged $D\left( r;r_{0} \right), R\left( \boldsymbol{T};\partial\Omega_{t} \right)$ and $E_{el}$ as a function of $\beta$**,** with fixed $\alpha={10}^{-2}$. $\beta$ controls the emphasis on the elastic energy during optimization, as well as the anti-aliasing. Hence, the converged $E_{el}$ decreases, the shape and the force field are smoothened with increasing $\beta$. The converged value of shape difference and traction increases strongly with increasing $\beta>{10}^{-3}$, while for $\beta<{10}^{-3}$ the converged values remain rather constant. Therefore, $\beta={10}^{-3}$ (and $\alpha={10}^{-2}$) is the best choice for the global optimization.

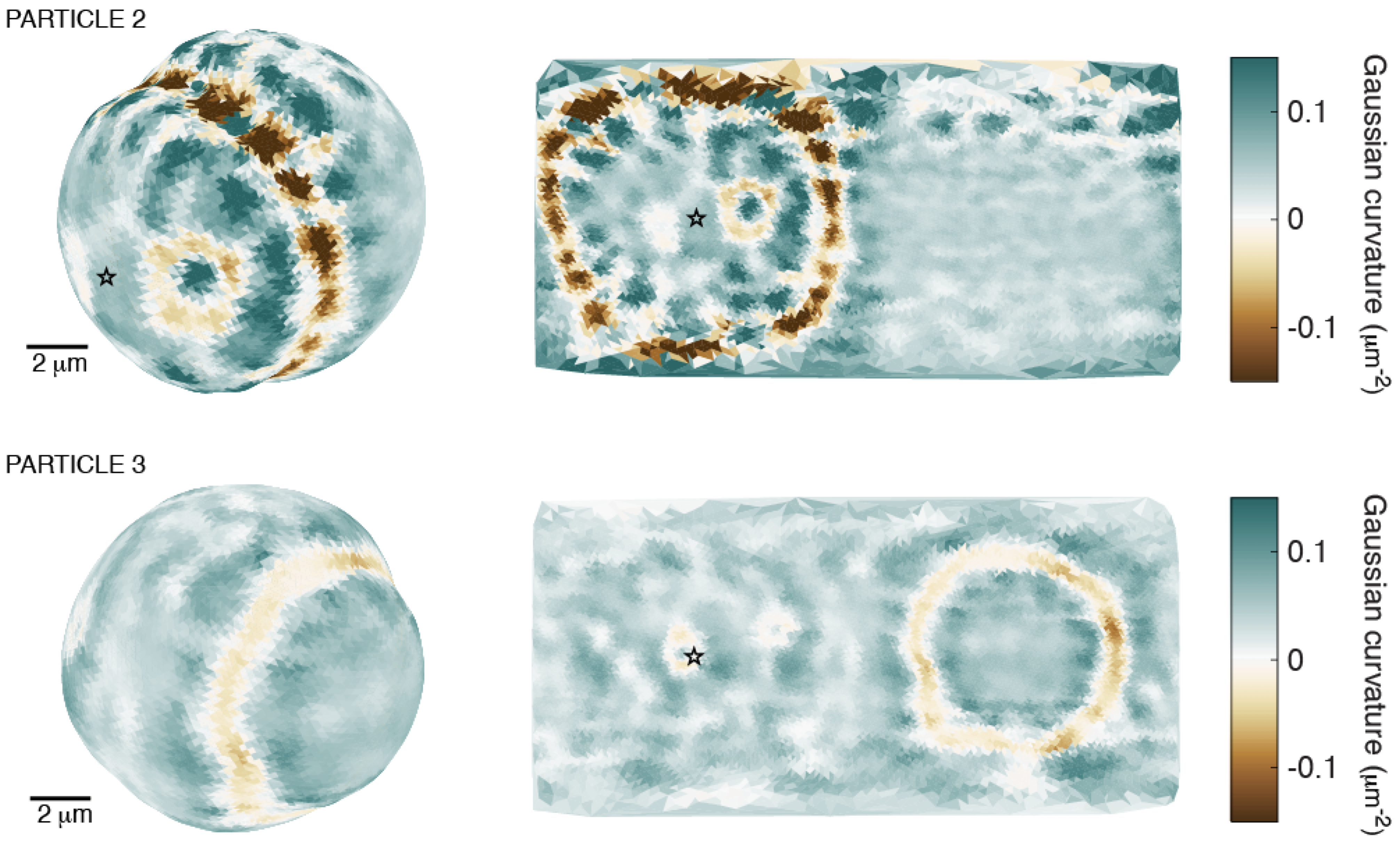

**Supplementary Figure 9: Gaussian curvature indicates ring homogeneity differences between particles.** A particle in the stage of pseudopod progression (top, particle 2, 34% engulfed, see also main text figure 4b) and a particle in the phase of cup closure (bottom, particle 3, 84% engulfed, see also main text figure 4c) are shown. Gaussian curvature is zero when one of the two principal curvatures is 0, positive when both principal curvatures have the same sign (a bulge or an indentation), and negative when the principal curvatures have opposite signs (a saddle point). This highlights homogeneities in the ring, which are striking in particle 2, but mostly absent in particle 3. Stars mark the phagocytic cup base. Left, projection of particle surface from a perspective that facilitates ring visualization. Right, equirectangular map projections (similar to figure 4 in the main text).

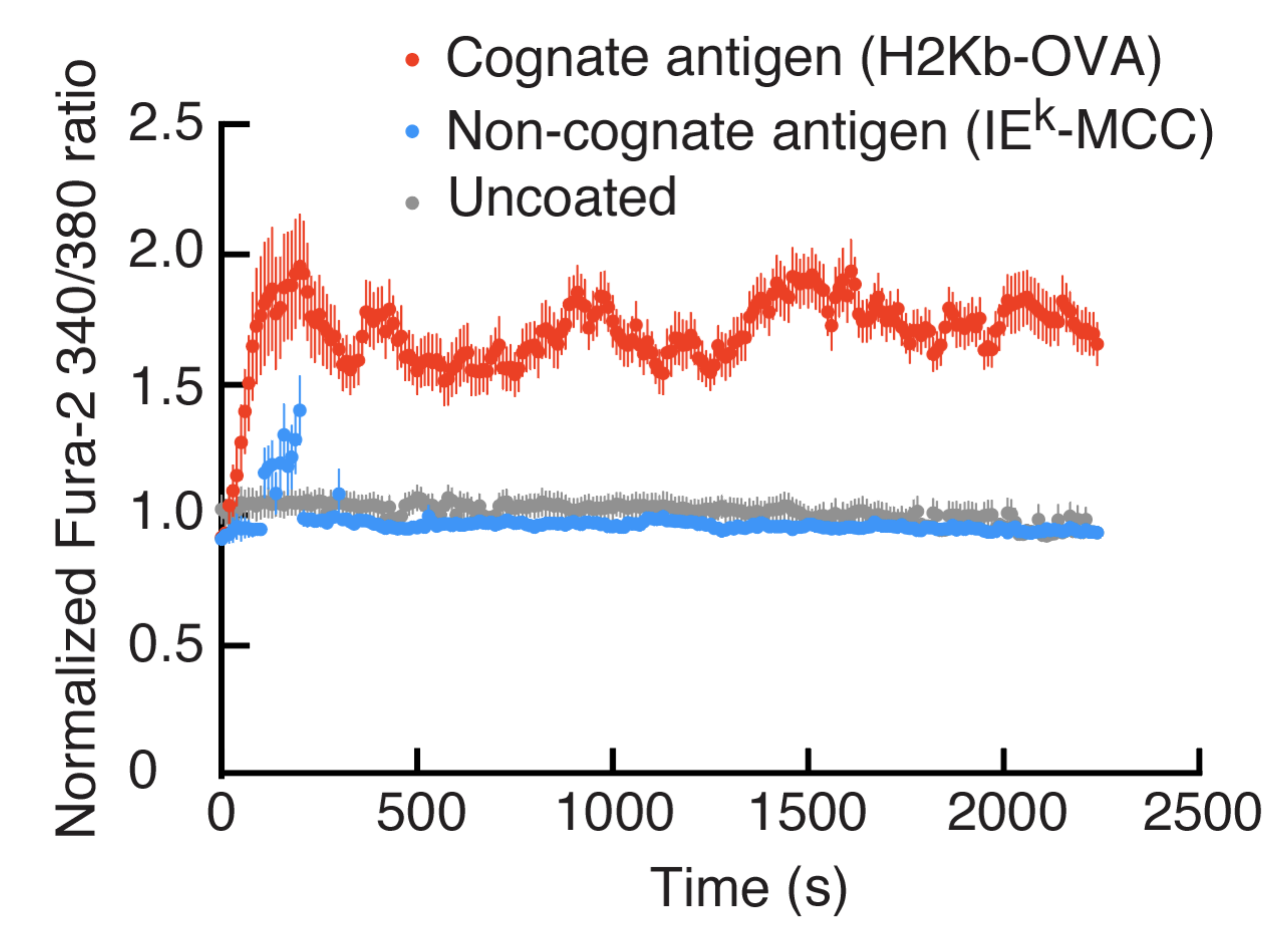

**Supplementary Figure 10: OT-I CA^2+^ response to DAAM-particles.** Day 7 CTLs from OT-I transgenic mice were loaded with the intracellular calcium (Ca^2+^) indicator Fura2-AM and incubated with DAAM-particles (8.6 μm, 4 kPa) coated with either cognate antigen (H2Kb-OVA and ICAM-1, n = 53 events) or non-cognate antigen (MCC-IEk and ICAM-1, n = 81 events), or left uncoated (n = 89 events). Normalized Fura ratios were measured by microscopy and graphed in time. Circles represent mean values and error lines represent standard error of the mean (s.e.m.). Data are representative of at least two independent experiments and are consistent across different stiffness regimes.

**Supplementary Table 1: Concentrations used for conjugation of the MPs**

| Particle | C_c_ | C BSA | C Ab1 | C Ab2 | C dye | Figure |
| --- | --- | --- | --- | --- | --- | --- |
|  | % (m/m) | (mg/mL) | (mg/mL) | (μg/mL) | mM |  |
| **Intraparticle homogeneity illustration** | | | | | | |
| DAAM | 2.3 | 5* | - | - | 2.32 | 1e |
| DAAM | 0.32 | 5* | - | - | 0.58 | S2 |
| **Lensing effect illustration** | | | | | | |
| DAAM | 0.32 | 19 | 0.09 | 4.5 | 0.58^§^ | 1i, S4 |
| PS | - | 0.04^†^ | 0.06 | 3 | - | 1i, S4 |
| **Mechanical characterization (AFM)/Determination resolution/Sphericity measurements** | | | | | | |
| DAAM | 2.3 | 77 | - | - | 2.32 | 1j, 1k, S5 |
| DAAM | 0.65 | 29 | - | - | 0.87 | 1k, S5 |
| DAAM | 0.32 | 19 | - | - | 0.58 | 1k, 2, S5 |
| **Measurements phagocytic efficiency** | | | | | | |
| DAAM | 2.3 | 77 | - | - | 0.58 | S7 |
| DAAM | 2.3 | 77 | 0.14 | - | 0.58 | S7 |
| **Phagocytic deformation assay** | | | | | | |
| DAAM | 0.32 | 19 | 0.09 | 4^‡^ | 0.58 | 3, 4, 5, S8 |

Concentration of crosslinker (C_c_), BSA, primary antibody (anti-BSA rabbit IgG), secondary antibody (anti-rabbit IgG) used for the various experiments. For all conjugations of deformable acrylamide-co-acrylic acid microparticles (DAAM-particles), the particle concentration was kept at 5% v/v. For polystyrene microparticles (PS MPs) particle concentration was 2% (v/v). BSA concentration was chosen such that enough BSA was present to bind 25% of the acrylic acid groups within the particles. *FITC-BSA instead of BSA was used for derivation, in which case a smaller amount of protein was used. ^†^For polystyrene particles a much lower concentration was used, since the protein cannot diffuse into the non-porous particle. In this case, 10x the amount to saturate the surface area of the particles was used. Primary antibody concentration was chosen such that enough IgG was present for opsonizing 2x the surface of the particles (although the primary antibody may diffuse into the particles). Secondary antibody concentration was picked such that enough antibody was present for ~0.1x surface saturation. ^‡^Immunostaining was performed after exposure of the beads to cells. Concentration of the carboxyl-reactive dye (TRITC-Cadaverine (Cad)) was chosen such that enough dye was present to bind 50% of the acrylic acid groups within the particles. ^§^Alexa Fluor 488-Cad was used instead of TRITC-Cad. Most right column indicates the figures for which each of these particles was used, where S indicates reference to supplementary figures.

**Supplementary Note 1: Spherical harmonic decomposition of particle shape**

For determining if there is a characteristic length scale of the deviations of non-deformed particles from a perfect sphere, the edge coordinates of adherent particles (those with the highest signal-to-noise ratio in figure 2a in the main text) were represented in Cartesian coordinates $\left( x_{i},y_{i},z_{i} \right), i=1,2,\ldots,n$. First, we determine the center $\left( X,Y,Z \right)$ and the radius $R$ of the particle by minimizing the sum of the squared distances between the data points to the spherical surface given by $\left( X,Y,Z,R \right)$:

$$\min_{\left( X,Y,Z,R \right)} \sum_{i} \left[ \left( x_{i}-X \right)^{2}+\left( y_{i}-Y \right)^{2}+\left( z_{i}-Z \right)^{2}-R^{2} \right]$$

The data points were then represented in spherical coordinates:

$$r_{i}=\sqrt{\left( x_{i}-X \right)^{2}+\left( y_{i}-Y \right)^{2}+\left( z_{i}-Z \right)^{2}},\theta_{i}=\arccos\frac{z_{i}-Z}{R},\varphi_{i}=atan2 \left( y_{i}-Y,x_{i}-X \right),$$

which can be treated as the radius of points being a function of co-latitude and longitude $r(\theta,\varphi)$. Considering the measurement error increases with the distance to the equator $(\theta=90^{\circ})$, the actual radius is corrected using weighted function $w\left( \theta\right)$ described in Supplementary Note 3:

$$\bar{r}_{i}\left( \theta_{i},\varphi_{i} \right)=w\left( \theta_{i} \right)\left( r_{i}-R \right)+R.$$

Since we only obtained data on the upper half of the particle, we copy and reverse the data points to form a complete sphere: $\bar{r}\left( \theta>90^{\circ},\varphi\right)=\bar{r}\left( 180^{\circ}-\theta,\varphi+\varphi_{0} \right)$, where $\varphi_{0}$ is a random angle between $\left( 0^{\circ},90^{\circ} \right)$ to avoid the symmetry of the upper and lower sphere. Next, using SHTools^2^ spherical harmonic analysis was performed on $\bar{r}\left( \theta,\varphi\right)$ using the least-square method:

$$\bar{r}\left( \theta,\varphi\right)=\sum_{l=0}^{l_{\max}} \sum_{m=-l}^{+l} \hat{r}_{l}^{m}Y_{l}^{m}\left( \theta,\varphi\right),$$

where $\hat{r}_{l}^{m}$ are the spherical harmonic coefficients, and $Y_{l}^{m}$ are the spherical harmonic functions defined in Supplementary Note 4. In the following analysis, the cutoff $l_{\max}=20$ was used to be consistent with the spherical harmonic method we use throughout the manuscript. Finally, the “per-$l$” power spectrum of the spherical harmonic coefficients are defined as

$$S_{l}=\sum_{m=-l}^{+l} \left( \hat{r}_{l}^{m} \right)^{2}$$

for presenting the magnitude of the coefficients as a function of spherical harmonic degree $l$.

**Supplementary Note 2: Dilation and smoothing of the edge of the traction-free surface**

For calculation of surface traction forces, the traction-free boundary region was dilated and its edge was smoothed. The rational for dilation is based on 1) lack of knowledge about the exact correspondence of the region of possible cellular force exertion on the target with the area of tight proximity between cell and target (where the fluorescent secondary antibody probe is excluded), 2) the notion that overestimation of the extent of the traction-free surface will likely have a more profound effects on the resultant forces than underestimation of this area. Smoothing of the edge was applied to prevent overestimation of high-frequency contributions in the force calculations.

Dilation by 1 μm was performed by excluding points from the traction free surface with great circle distance <1 μm to any point part of the edge of the traction-free boundary. Then a weight function is defined as:

$$w_{t}\left( \theta,\varphi\right)=\left\{ \begin{aligned} 1, beyond the dilated edge \left( completely traction free \right) \\ \left( 0,1 \right), between non\text{-}dilated and dilated edge \left( smoothed edge region \right) \\ 0, inside the non\text{-}dilated region \left( completely in contact \right) \end{aligned} \right.$$

In between the smoothed edge region, we smoothly interpolate the $w_{t}$ values between 0 and 1. In particular, we constructed $w_{t}\left( \theta,\varphi\right)$ as following: we first assign 0.5 to the values on the transitional region:

$$w_{t}^{'}\left( \theta,\varphi\right)=\left\{ \begin{aligned} 1, beyond the dilated edge \left( completely traction free \right) \\ 0.5, between non\text{-}dilated and dilated edge \left( smoothed edge region \right) \\ 0, inside the non\text{-}dilated region \left( completely in contact \right) \end{aligned} \right.$$

Then a spherical harmonic expansion is applied to $w_{t}^{'}\left( \theta,\varphi\right)$:

$$w_{t}^{'}\left( \theta,\varphi\right)=\sum_{l,m} \hat{w}_{t,lm}^{'}Y_{l}^{m}\left( \theta,\varphi\right),$$

and the edge-smoothing is applied to $w_{t}$ by damping the high frequency coefficients:

$$w_{t}\left( \theta,\varphi\right)=\sum_{l,m} \hat{w}_{t,lm}^{'}q\left( l \right) Y_{l}^{m}\left( \theta,\varphi\right),$$

where

$$q\left( l \right)=\left\{ \begin{aligned} 0, l>l_{hi} \\ \sin^{2} \left( 2\pi\frac{l_{hi}-l}{l_{hi}-l_{lo}} \right), l_{lo}\leq l\leq l_{hi} \\ 1, l<l_{lo} \end{aligned} \right.$$

which cuts off the coefficients smoothly between $l_{hi}$ and $l_{lo}$. In this paper, we use $l_{lo}=12$ and $l_{hi}=20.$

Finally, the corrected residual traction magnitude is calculated as:

$$R\left( \boldsymbol{T};\partial\Omega_{t} \right)=\sqrt{\frac{1}{m}\sum_{i} \left\| \left[ \boldsymbol{T}_{i}\left( \theta_{i}^{s},\varphi_{i}^{s} \right)\cdot\boldsymbol{w}_{n} \right] w\left( \theta_{i}^{s} \right) w_{t}\left( \theta_{i}^{d},\varphi_{i}^{d} \right) \right\|^{2}}$$

The traction in the contact region is hence not considered in$R\left( \boldsymbol{T};\partial\Omega_{t} \right)$, while the traction on the completely traction-free region is fully considered and the traction on the edge is partially considered.

**Supplementary Note 3: Spherical Harmonics definition**

The definition of spherical harmonic functions in this paper is as following:

$$Y_{l}^{m}\left( \theta,\varphi\right)=\left\{ \begin{aligned} \bar{P}_{l}^{m}\left( \cos\theta\right) e^{im\varphi}, m\geq0 \\ \left( -1 \right)^{\left| m \right|}\left[ Y_{l}^{\left| m \right|} \right]^{*},m<0 \end{aligned} \right.,$$

where $\left[ Y_{l}^{\left| m \right|} \right]^{*}$is the complex conjugate of $Y_{l}^{\left| m \right|}$, and $P_{l}^{m}\left( \mu\right)$ is the normalized associated Legendre function with the complex $4\pi$-normalization which is defined as following:

$$\bar{P}_{l}^{m}\left( \mu\right)=\sqrt{\left( 2l+1 \right)\frac{\left( l-m \right)!}{\left( l+m \right)!}}P_{lm}\left( \mu\right)$$

The unnormalized associated Legendre functions are:

$$P_{lm}\left( \mu\right)=\left( 1-\mu^{2} \right)^{\frac{m}{2}}\frac{d^{m}}{d\mu^{m}}P_{l}\left( \mu\right),$$

where the Legendre function $P_{l}\left( \mu\right)$ is defined in Supplementary Note 2. Spherical harmonic functions are orthogonal for different $\left( l,m \right)$ degrees:

$$\iint_{\partial\Omega} \left[ Y_{l}^{m}\left( \theta,\varphi\right) \right]^{*} Y_{l^{'}}^{m^{'}}\left( \theta,\varphi\right) dS=4\pi\delta_{ll^{'}}\delta_{mm^{'}},$$

where $\delta_{ll^{'}}$ is the Kronecker delta function.

Given the orthogonality property of spherical harmonic functions, it is straightforward to evaluate the elastic energy by convolving displacement field and traction field on the spherical surface, which equals to the inner product of spherical harmonic coefficients:

$$E_{el}=\frac{1}{2}\iint_{\partial\Omega} u_{j}\left( \theta,\varphi\right)\cdot T_{j}\left( \theta,\varphi\right) dS=2\pi\sum_{j,l,m} \left[ \hat{u}_{jlm} \right]^{*}\hat{T}_{jlm}$$

where $\left[ \hat{u}_{jlm} \right]^{*}$ are the complex conjugates of the coefficients $\hat{u}_{jlm}$.

**Supplementary Note 4: Gauss-Legendre quadratic (GLQ) meshing, latitude and resolution weighing function**

Gauss-Legendre quadratic mesh with degree $L$ is exact if the sampling function $f\left( \theta,\varphi\right)$ is terminated within $L$ degrees of spherical harmonics. The longitude nodes are uniform on longitude with $360^{\circ}/\left( 2L+1 \right)$, and the latitude nodes correspond to zeros of the Legendre Polynomial of degree $l=L+1$:

$$P_{l}\left( \mu\right)=\frac{1}{2^{l}l!}\frac{d^{l}}{d\mu^{l}}\left( \mu^{2}-1 \right)^{l},$$

where $\mu=\cos\theta$. In total, GLQ has $n_{\mathrm{node}}=\left( L+1 \right)\times\left( 2L+1 \right)$ nodes on $\left( \theta,\varphi\right)$ space.

GLQ meshing introduces a higher node density at the poles of the sphere than at the equator. To avoid overrepresentation of the contribution of the shape and traction at the poles, the nodes are weighted by weight function $w\left( \theta\right)=\sin\theta$ during the evaluation of the shape difference$D\left( r;r_{0} \right)$ and the traction magnitude $R\left( \boldsymbol{T};\partial\Omega_{t} \right)$. In addition, we also include the weighing vector $\boldsymbol{w}_{n}=\left[ 1,1,1/3 \right]^{T}$in Cartesian coordinates to address that the measurement noise level in the axial-direction is higher than in the lateral directions.

$$D\left( r;r_{0} \right)=\sqrt{\frac{1}{n}\sum_{i} \left\| \left[ \left( r_{i}^{d}-r_{0}\left( \theta_{i}^{d},\varphi_{i}^{d} \right) \right)\hat{\boldsymbol{r}}\left( \theta_{i}^{d},\varphi_{i}^{d} \right)\cdot\boldsymbol{w}_{n} \right] w\left( \theta_{i}^{s} \right) \right\|^{2}}$$

$$R\left( \boldsymbol{T};\partial\Omega_{t} \right)=\sqrt{\frac{1}{m}\sum_{\left( \theta_{i}^{d},\varphi_{i}^{d} \right)\in\partial\Omega_{t}} \left\| \left[ \boldsymbol{T}_{i}\left( \theta_{i}^{s},\varphi_{i}^{s} \right)\cdot\boldsymbol{w}_{n} \right] w\left( \theta_{i}^{s} \right) \right\|^{2}}$$

**SUPPLEMENTARY DISCUSSION**

**Superresolving the particle boundary by edge localization microscopy (ELM)**

In the radial direction, we have been able to resolve the particle boundary with accuracy beyond the diffraction limit (d_x,y_ ≈ 200 nm, d_z_≈ 600 nm). Such resolution is accomplished similarly to single-molecule localization techniques^3,4^; the particle edge, representing an optically distinct feature, can be localized with precision that increases approximately with 1/√N, where N is the number of photons collected. For a homogeneously labeled particle, the radial intensity profile can be approximated as the convolution of a step function, representing the homogeneous and dense presence of fluorophores, with a Gaussian function representing the emission spot, *i.e.* the point spread function (PSF), of a single fluorophore. The derivative of this convolution is, in fact, a Gaussian itself, localized at the particle edge. Hence, edge superlocalization can be performed by fitting a Gaussian to the derivative of the intensity profile at the particle edge, where the discrete derivative can, for example, be approximated using the 3D Sobel operator^5^.

**Synchronized particle polymerization using oil-soluble polymerization initiators**

The particle-to-particle variation we observe in optical and mechanical properties is remarkably low. For MPs with a refractive index ~1.4, previously a spread on the order of 0.001 was reported^16^, while we observe a ~10 fold lower spread of ~0.0001. This homogeneity, which is also present in mechanical properties (CV = 0.08 - 0.30 for stiff to soft particles), may be related to our choice for an oil-soluble polymerizing agent. Water-soluble initiators, such as ammoniumpersulfate (APS) have frequently been used for polyacrylamide polymerization, and were also used previously for particle synthesis. Such initiators must be present in the gel mixture during emulsification, since they are not oil-soluble. It was previously shown that use of such initiators can lead to differential particle properties (optical and mechanical) over the extrusion period.^6^ This could be attributed to polymerization of the gel mixture before extrusion, which occurs, albeit slowly without TEMED, over the relatively long extrusion period (**≳**1h).^6^ Another potential explanation is based on the rapid degradation of APS in aqueous solutions, which could result in a higher concentration of APS and more efficient polymerization of droplets extruded early on, while droplets extruded later will polymerize less efficiently and will therefore have different properties. Regardless of the exact mechanisms that underlie the particle property dependence on extrusion time, use of an oil-soluble initiatior (*e.g.* AIBN) prevents such problems as it prevents polymerization of the gel mixture before extrusion and initiator degradation, and allows triggering polymerization in a entirely synchronized fashion. Using an oil-soluble initiator therefore likely improves particle-to-particle homogeneity. Finally, the use of AIBN did not seem to affect the uniformity of the particle polymer network, as evidenced from the homogeneous radial fluorescent staining of the particles (Fig. 1e, Supplementary Fig. 2), which indicates that AIBN diffusion into the microdroplets did not result in any radial particle inhomogeneities.
